# Multi-objective management of naturally regenerating beech forests – An ecological-economic optimization approach

**DOI:** 10.1101/2024.06.07.597908

**Authors:** Markus E. Schorn, Martin F. Quaas, Hanna Schenk, Christian Wirth, Nadja Rüger

## Abstract

How can we meet economic objectives of timber harvesting while maintaining the functioning of diverse forest ecosystems? Existing forest models that address this type of question are often complex, data-intensive, challenging to couple with economic optimization models, or can not easily be generalised for uneven-aged mixed-species forests. Here, we develop an ecological-economic optimization model, which integrates a state-of-the-art demographic forest model with a continuous cover forestry harvesting model to optimise efficient and sustainable timber harvesting. As a proof-of-concept, we apply the model to a beech-dominated forest in the Hainich-Dün region in Thuringia, Germany, with the goal of optimising multiple objectives such as timber yield and the biodiversity value of the forest. The ecological module is the Perfect Plasticity Approximation (PPA) demographic forest model that simulates forest dynamics based on individual tree growth and survival rates in the canopy and understory layers, respectively, as well as recruitment rates. We used repeated forest inventory data from a 28-ha forest plot to quantify these demographic rates and validated the predictions of the ecological module against the structure of old-growth beech forests in Europe. The economic module includes the optimization of net revenues (market revenues net of harvesting cost) from harvesting timber. As an indicator of the biodiversity value of the forest, we use the number of retained habitat trees (>70 cm diameter). The forest model delivered reasonable predictions of structural attributes of unmanaged old-growth beech forests. When net revenues from timber harvest were maximised, trees were logged when they reached 55 cm in diameter. This is similar to current management practices in beech forests. We found a linear trade-off between timber net revenues and biodiversity value with about 2.5% of the maximum benefit of timber harvest being lost with each additionally retained habitat tree. We established a generic ecological-economic modelling framework that reliably represents forest dynamics as well as optimising forest management. To our knowledge, this is the first forest model for central European forests capable of identifying optimal harvesting over the full set of feasible strategies, rather than merely comparing predefined management scenarios. The framework can be extended to mixed-species forests and support forest management for diverse ecosystem services.

## 1. Introduction

Forests face intense and diverse societal demands and interests. Private forest ecosystem services such as timber production for construction or biomass production for energy supply and industrial uses are of high economic relevance, especially in rural regions (BMEL, 2024). In addition, forests provide multiple common-good services which are gaining in importance (Biber et al., 2015; Bugmann et al., 2017; Baeten et al., 2019; IPBES, 2019; FAO, 2022). These include the provision of habitat for a large fraction of biodiversity (FAO, 2022), the sequestration and storage of carbon (Riedel et al., 2019; Beudert and Leibl, 2020; FAO, 2022), or functional resilience against climate extremes (Mahecha et al., 2022), among others. There is plenty of evidence that forests with a heterogeneous size structure, i.e. uneven-aged forests, and forests with a diverse range of tree species provide higher levels of multiple ecosystem services than even-aged monocultures (e.g. Gamfeldt et al., 2013; van der Plas et al., 2016; Felipe-Lucia et al., 2018; Pardos et al., 2021; Messier et al., 2022). However, scientific tools that allow optimising the management of these forests for multiple purposes are scarce.

Individual-based forest dynamics models, such as SILVA, BWinPro, Sybila, or MOSES (Pretzsch, 2019) allow simulating detailed management scenarios of uneven-aged and mixed-species forests (Söderbergh and Ledermann, 2003). However, their parameterization is data- and labour-intensive and, due to their complexity, they cannot directly be coupled with economic optimization models, and their results are not easily generalizable across regions or tree species. Thus, a truly multi-objective optimization of forest management has largely been restricted to monocultures (e.g. Assmuth et al., 2018), to uneven-aged and mixed-species forests in the boreal zone (e.g., Tahvonen et al., 2019), or to the comparison of pre-defined management scenarios (Augustynczik et al., 2020; Gutsch et al., 2018). For temperate forests, it is still in its infancy, due to the complexity of the associated silvicultural forest simulation models (Lafond et al., 2017). Therefore, forest models are needed that meet the requirements of economic optimization and combine sufficient ecological realism to reliably simulate the dynamics of managed uneven-aged mixed-species forest stands with easy transferability to other forests of interest.

Here we propose that the Perfect Plasticity Approximation (PPA) model (Purves et al., 2008; Strigul et al., 2008) is a good candidate to meet these requirements. The PPA model simulates the dynamics of uneven-aged mixed-species forest stands and directly incorporates competition for light by simulating two (or more) dynamic canopy layers. The PPA model is straightforward to parameterize because it is based on a small set of species-specific demographic rates (growth and survival in canopy and understory layer respectively, recruitment) which can directly be derived from forest inventory data, and has been shown to produce reliable predictions of long-term forest dynamics in temperate and tropical forests (Francis et al., 2023; Purves et al., 2008; Rüger et al., 2020). The PPA model is analytically tractable because the assumption of perfect plasticity (i.e., each tree can place its crown area anywhere in the horizontal plane; Strigul et al., 2008) enables the individual-based model to be recast as a demographic model, tracking the fate of cohorts rather than individuals.

For economic optimization, we transform the cohort model into a size class model with parameters that are calculated from the demographic rates of the cohort model, and numerically optimise size-dependent harvesting for profit from timber yield assuming constant marginal harvesting costs and timber prices. As a proof-of-concept, we apply the model to an uneven-aged beech-dominated forest in the Hainich area, Germany (Huss and Butler-Manning, 2006). To validate the ecological submodel, we compare simulated structural characteristics of an unmanaged old-growth forest with information on forest structure in European old-growth beech forests. To validate the economic submodel, we compare optimal timber yield predicted by the model to observed yields in uneven-aged beech forests in Germany. Additionally, we also maximise the harvested timber volume under the constraint of retaining a given number of large habitat trees (>70 cm diameter) for biodiversity conservation to identify the trade-off between these ecosystem services. The number of large old trees is an important indicator of the biodiversity conservation value of a forest because these trees provide a high number of microhabitats (Gao et al., 2015; Lindenmayer, 2017; Mergner, 2021; Prevedello et al., 2018). While habitat tree retention is recommended by forestry authorities in several federal states in Germany (e.g., ForstBW, 2010; Niedersächsische Landesforsten, 2018; Hessen-Forst, 2022) and required for some certifications (FSC, 2023; FSC Deutschland, 2018; PEFC Deutschland, 2020), the economic effects are rarely discussed, especially for continuous-cover forestry (Gustafsson et al., 2020).

In summary, we propose an ecological-economic optimization framework for European beech forests as a methodological basis to support forest management decisions integrating economic and conservation goals. In this proof-of-concept, we test whether the ecological model reliably predicts forest dynamics and whether the timber yield optimization output provides realistic target diameters for selection forestry. We also evaluate the trade-off between timber yield and biodiversity conservation value of the forest. Because the forest model relies on few demographic parameters that can easily be estimated from forest inventory data, this framework provides a unique basis for the development of widely applicable ecological-economic forestry models. It can be extended to mixed-species forests and to the optimization of multiple services such as biodiversity conservation (as shown here) or carbon storage.

## 2. Methods and materials

We propose a novel ecological-economic model optimization framework based on the Perfect Plasticity Approximation model (PPA, Purves et al., 2008). The PPA model is a cohort model that is continuous in tree diameter. For economic optimization, we transform the cohort model into a size class model that has discrete size classes. We parameterize both models for a beech-dominated forest in the Hainich National Park, Germany, and validate the ability of the size class model to approximate the cohort model and the structure of unmanaged old-growth forests. We use the size class model to optimise harvesting for timber yield with and without a constraint on the number of large habitat trees to be retained in the forest as a proxy for biodiversity conservation.

### 2.1 The cohort model

The PPA model simulates forest dynamics based on a small set of demographic rates (growth, mortality, and recruitment) and accounts for height-structured competition for light by distinguishing two dynamic discrete canopy layers: the canopy layer, where trees have access to light and the understory layer, where trees are shaded. Trees are assigned to the canopy layers based on their size and the size of their neighbours. The tallest trees are assigned to the top canopy layer as long as their cumulative crown area does not exceed the simulation area. Canopy gaps are filled by the tallest trees from lower canopy layers without regard for their horizontal position (perfect plasticity assumption, Strigul et al., 2008). Trees in the two canopy layers are assigned different growth and mortality rates. Tree species typically have higher diameter growth rates and lower mortality rates in the canopy layer (*g*_*C*_, *μ*_*C*_) than in the understory layer (*g*_*U*_, *μ*_*U*_). The recruitment of new trees *R* is a constant rate per year and ha (Table 1, Rüger et al., 2020). Details of the cohort model can be found in the original publications (Purves et al., 2008; Strigul et al., 2008).

**Table 1.**
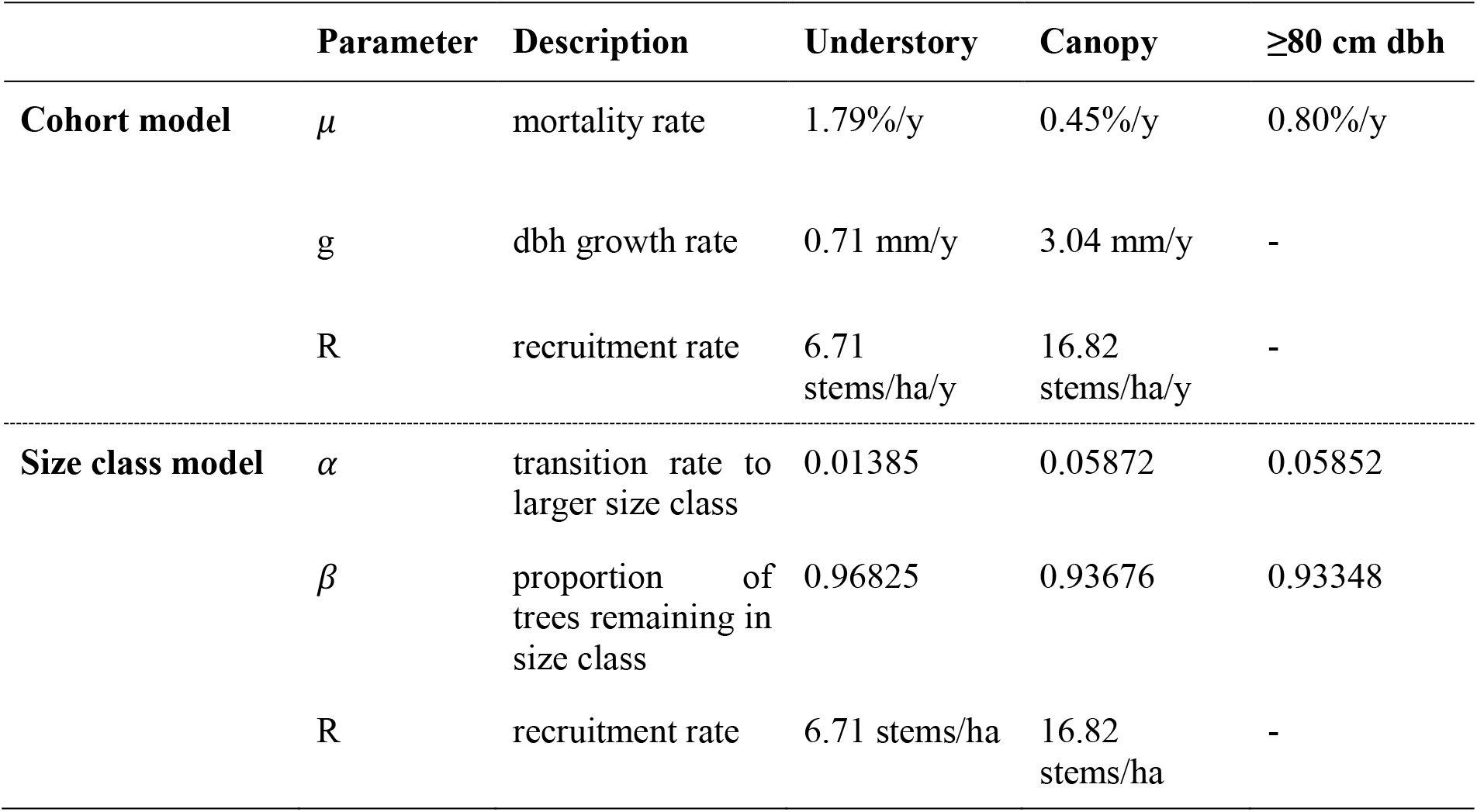
Model parameters for the cohort and size class models for trees in light (Canopy) and in shade (Understory), as well as for very large trees (≥80 cm dbh). Recruitment rates R refer to the number of trees growing over 1.3 m height per year and hectare in situations of an open canopy (Canopy) or a closed canopy (Understory).

Here, we deviate from previous model versions in two aspects. First, we introduce elevated mortality rates for very large canopy trees (Francis et al., 2023), based on the findings of Holzwarth et al. (2013) who observed U-shaped mortality curves at the same site. Second, we make the recruitment rate dependent on canopy openness, such that we introduce a recruitment rate for situations when the canopy is or is not fully closed (total crown area ≥1 or <1 m^2^/m^2^, respectively).

### 2.2. The size class model

For the purpose of optimization, we transformed the model to a transition model with discrete size classes *s*. Discrete size classes are increasingly standard in economic-ecological optimization models, as they allow an optimization over the full set of possible solutions (in particular, uneven-aged forest management), while still maintaining numerical tractability (Tahvonen, 2015). The number of trees that are promoted to the next size class in each time step is determined by a compound ‘growth-and-survival rate’ *α* and the number of trees remaining in the same size class is determined by a ‘survival rate’ *β*. Both transition rates *α* And *β* are defined for size classes that are in the canopy and size classes that are in the understory and depend on the growth and mortality rates of canopy and understory trees derived from forest inventory data. Additionally, α and β are defined for very large canopy trees (≥80 cm diameter at breast height, dbh):

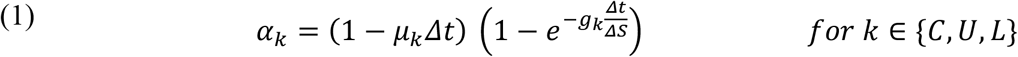

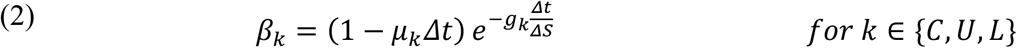

where *μ*_*C*_ and *μ*_*U*_ are the mortality rates of canopy and understory trees, respectively, *g*_*C*_ and *g*_*U*_ are the growth rates of canopy and understory trees, respectively, *μ*_*L*_ and *g*_*L*_ are mortality and growth rates of very large canopy trees (≥80 cm dbh), and *ΔS* is the width of a size class in cm.

*α* and *β* of a size class *s* depend on the total crown area of all trees in bigger size classes, i.e. the shading from above normalised to the simulation area *A*_*s*_ (i.e. in m^2^/m^2^). Using a smoothing factor *σ* = 0.01, *α(A*_*s*_) and *β(A*_*s*_) approximate *α*_*C*_ or *β*_*C*_ for *A*_*s*_<1 (i.e. open canopy), *α*_*U*_ or *β*_*U*_ for *A*_*s*_>1 (i.e. closed canopy), and *α*_*L*_ or *β*_*L*_ for trees ≥80 cm dbh (see Appendix A, Fig. A1):

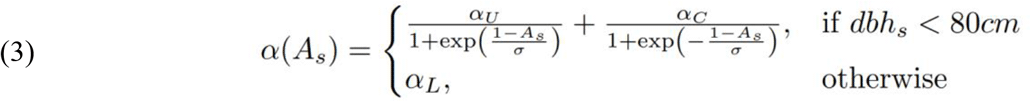

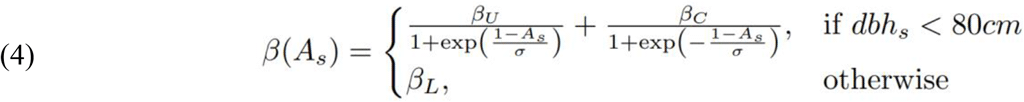

where *dbh*_*s*_ is the dbh of a tree of size class s (= *s* · *ΔS*) and *A*_*s*_ is given by:

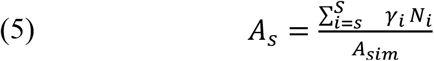

where *γ*_*s*_ is the crown area of a single tree of size *s, N* is the number of trees in size class *s* and *A*_*sim*_ is the simulated area. The allometric diameter-crown area relationship used to calculate *γ*_*s*_ is given in Appendix B.

The number of trees *N*_*st*_ in size class s = 1, 2, …, S and period t = 1, 2, … is then given by:

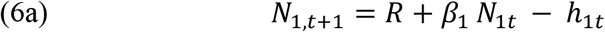

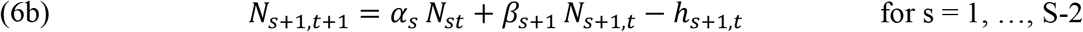

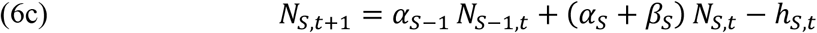

where *h*_*st*_ is the number of harvested trees per ha per size class and the recruitment rate *R* depends on shading (*R*_*C*_ for *A*_*st*_<1 and *R*_*U*_ for *A*_*st*_>=1, Table 1).

The width of the size classes *ΔS* and the time step Δt for the calculation of the transition rates α and β should be selected such that, based on the annual growth rate, individuals can not skip a size class. We use S = 30 size classes of width *ΔS* = 5 cm, resulting in an intrinsic maximum dbh of 150 cm. We use a time step of Δt = 1 year to calculate the transition rates α and β. We simulate an area of *A*_*sim*_ = 1 ha.

### 2.3 Model parameterization

#### 2.3.1 Study site and forest inventory data

We used data from a 28.5 ha plot of mature deciduous forest (‘Weberstedter Holz’; 51°06′ N, 10°31′ E) located in the Hainich National Park, Thuringia, Germany, to derive the demographic rates for the cohort model. The climate at the study site is suboceanic/subcontinental with a mean annual temperature around 7.5–8°C and mean annual precipitation around 750–800 mm (Holzwarth et al., 2013). Altitude is approximately 440 m a.s.l. with a gentle north facing slope (Huss and Butler-Manning, 2006). Soils at the study area are generally Luvisols developed from Pleistocene loess over Triassic limestone. The stand was managed as a coppice-with-standards until about 1890, after which it was gradually transformed into a beech selection forest. After 1965, when it became part of a military training ground, no regular forest management was applied apart from cutting a few single high-value trees. Management ceased completely in 1997, when the area was declared a core zone of the Hainich National Park (Butler-Manning, 2007; Mund, 2004). The main tree species at the study site is European beech (*Fagus sylvatica* L.) along with ash (*Fraxinus excelsior* L.), hornbeam (*Carpinus betulus* L.), sycamore (*Acer pseudoplatanus* L.) and wych elm (*Ulmus glabra* Huds.). Beech accounts for 89.8% of the stems >1.3 m height and 67.2% of the basal area. The diameter at breast height (1.3 m, dbh) as well as the exact location of all trees taller than 1.3 m were measured, marked, and re-measured in 1999, 2007, and 2013.

#### 2.3.2 Demographic rates

From this data, we derived growth and mortality rates for European beech for canopy and understory trees >1.3 m height (i.e. 0 cm dbh). We assigned trees to either the canopy or the understory based on their size and the size of their neighbours following the PPA approach (Purves et al., 2008). To do this, we estimated the height and crown area of all trees using species-specific allometric equations that we derived from data from 1,316 trees (Appendix B; Fleck et al., 2011; Holzwarth et al., 2015). We then divided the plot into quadratic subplots of 400 m^2^ in size. As a result, we neglected data from 3.3 ha in the edge regions of the plot because of its non-rectangular shape. Next, we sorted trees by height within subplots. Starting from the largest, we assigned trees to the canopy layer as long as the cumulative estimated area of their crowns did not exceed the subplot area. Smaller trees were assigned to the understory. Due to the uncertainty associated with the canopy layer classification, we only used trees that were assigned to a given canopy layer with some certainty for the calculation of demographic rates. We only used trees for the calculation of canopy and understory layer rates, where larger trees had <=0.8 or >1.2 m^2^/m^2^ crown area. We used data from all trees for the canopy layer classification but only calculated demographic rates for beech, because in this proof-of-concept, we focus on a single-species model for maximum simplicity.

We calculated individual annual absolute dbh growth for all beech trees in both census intervals, 1999–2007 and 2007–2013. We removed five growth outliers (>5 cm/year). We then calculated species-level growth rates per canopy layer as the mean growth of all beech individuals across census intervals (Table 1). We determined annual mortality rates per canopy layer (*μ*_*k*_) as

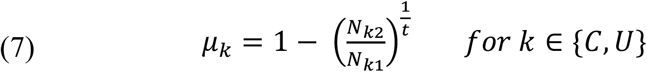

with *N*_*k*1_ being the number of living individuals at the beginning of the census interval, *N*_*k*2_ being the number of individuals remaining alive at the end of the census interval and *t* being the mean census interval length in years (i.e. 7 years, Table 1).

To determine recruitment rates, we first calculated the number of recruits above the 1.3 m height threshold per ha and year for each of the 400-m^2^ subplots separately per census interval. Next, we determined the shading of each subplot as the total crown area at the beginning of the census interval divided by the subplot area. The closed-canopy recruitment rate RU is the mean of recruitment rates across subplots and census intervals with shading ≥1 and the open-canopy recruitment rate RC is the mean across subplots and census intervals with shading <1 (see Appendix C, Fig. C1).

### 2.4 Model validation

To test the ability of the size class model to approximate the cohort model and the structure of unmanaged forests, we determined basal area, stem density, density of very large trees (≥80 cm dbh), maximum dbh, aboveground biomass, standing timber volume, as well as size distributions in equilibrium in the size class model and the cohort model. To determine equilibrium structure in the size class model, we fix the harvesting rate to 0. The cohort model oscillates in equilibrium (Appendix D, Fig. D1). Therefore, we average all structural variables over one oscillation period (325 years, i.e. from 1675 years to 2000 years). The maximum dbh in the cohort model is derived as the dbh at which stem density falls below 0.0398 stems/ha, which equals 1 stem in the observed area at the Weberstedter Holz (25.2 ha). We compare predicted forest structure with observations from the study site as well as from unmanaged beech-dominated forests in Europe (Vandekerkhove et al., 2018; Stillhard et al., 2019; Nagel et al., 2023; Appendix E, Table E1).

### 2.5 Optimization of harvesting

We are interested in optimal long-term steady states. As we focus on sustainable forest management, we ignore economic discounting, and we also focus on uneven-aged forest management. Accordingly, the forest structure subject to harvesting is characterised by model (6a-c) for the case when number of trees *N*_*s*_ in each size class s = 1, 2, …, S remains constant over time.

We maximise the net revenues from timber harvesting across all size classes at equilibrium:

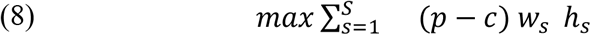

with *h*_*s*_ denoting the number of harvested trees per size class (see Equations 6a-c), *p* denoting the wood price per m^3^, *c* denoting harvesting costs per m^3^, and *w*_*s*_ denoting the merchantable timber volume in m^3^ of a single tree of size class *s* (Döbbeler et al., 2006; Hessenmöller et al., 2018):

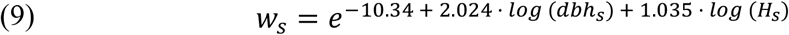

where *H*_*s*_ is the height in m of a tree of size class *s* (Appendix B):

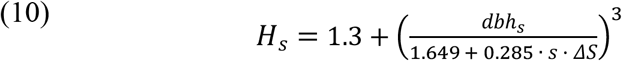

and *dbh*_*s*_ is the diameter breast height of a tree of size class s (= *s* · *ΔS*).

Wood prices fluctuate between years and quality grades. Here, we use an average figure of 65 Euros/m^3^ of stem wood for beech (Förster, 2011). In line with the recommendations for private forest owners, we assume that the harvesting costs *c* are computed per cubic metre of harvested merchantable timber. A typical figure is that costs are about 25% of revenues (Förster, 2011), yielding net revenues of 48.75 Euros/m^3^.

We compare net-revenues-maximising harvest with observations from managed uneven-aged beech forests in the study area (Hessenmöller et al., 2018; Schall et al., 2018). Since natural mortality rates in managed forests are usually lower than in unmanaged forests (Meyer et al., 2022; Schall et al., 2018), we also run the optimization with a reduced canopy mortality rate of 0.1%.

We are furthermore interested in the possible trade-off between private benefits from timber harvesting and non-economic values derived from a biodiverse forest ecosystem. To optimise the harvest while accounting for the value of large habitat trees for biodiversity conservation, we add the constraint of retaining the largest n trees ≥70 cm dbh per hectare until their natural death. We iterate over different values of n to find the Pareto front representing the trade-off between maximum harvest and biodiversity conservation value of the forest.

For numerical optimization, we use the state-of-the-art interior point algorithm implemented in AMPL with Knitro (Byrd et al., 1999, 2006). The multi-start optimizations start with 10,000 random initial conditions when only maximising yield and with 100 different initial conditions for each value of *n* when including the habitat trees constraint. Initial conditions in the second step are normally distributed around the optimal values from the first step using a standard deviation of 1% of the mean.

## 3. Results

### 3.1 Validation of equilibrium structure in unmanaged forests

The predicted structure in non-managed equilibrium forests compares favourably with observations in European non-managed beech-dominated forests for the cohort and the size class model (**Figure 1**, Appendix E, Table E1). The cohort model predicts a stand basal area of 45.4 m^2^/ha (of which 43.4 m^2^/ha are from trees ≥30 cm dbh), aboveground biomass of 556 t/ha, a total stem density of 480 stems/ha, and a density of very large trees (≥80 cm dbh) of 27.2 stems/ha at equilibrium. The size class model predicts a basal area of 44.4 m^2^/ha (of which 41.9 m^2^/ha from trees ≥30 cm dbh), aboveground biomass of 530 t/ha, a total stem density of 446 stems/ha, and a density of very large trees (≥80 cm dbh) of 30.1 stems/ha. Hence, model predictions lie at the upper range of observed values. However, some of the forests still recover from previous management and gain in basal area, number of large trees, and maximum observed diameter (e.g. Kersselaerspleyn, Razula, Weberstedter Holz; **Figure 1**, Appendix E, Table E1). The structure of the pristine forest in Uholka is very similar to our model predictions. While the size class model has an intrinsic maximum diameter of, in our case, 150 cm, the maximum diameter in the cohort model is larger than the largest observed diameter (**Figure 1B**). Overall, the size class model approximates the equilibrium forest structure of the cohort model well (**Figure 2**).

**Figure 1:**
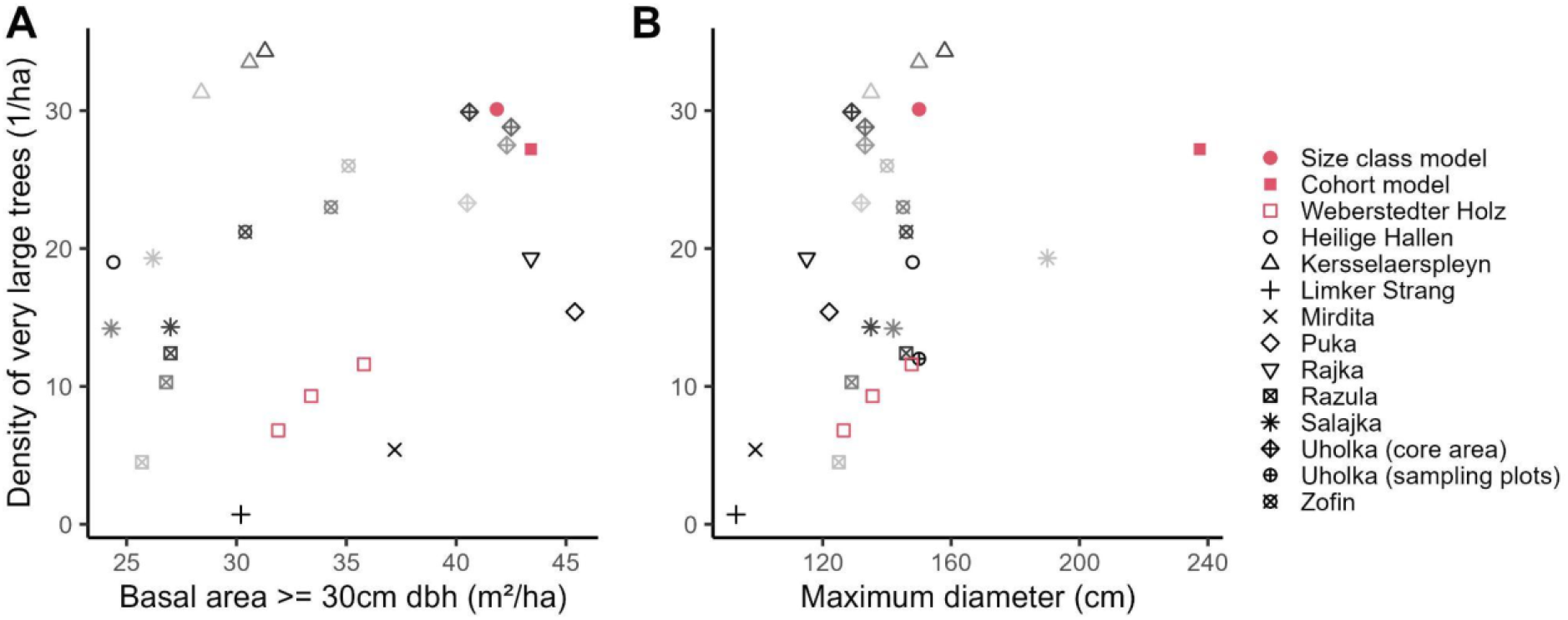
Basal area, density of very large trees (≥80 cm dbh) (A) and maximum diameter (B) of the model outputs at equilibrium compared with values from Weberstedter Holz as well as literature values. Red points represent the model outputs, grey and black points represent reference data from currently unmanaged beech-dominated forests in Central Europe (Vandekerkhove et al., 2018; Nagel et al., 2023, Table D1). Point shapes represent different forest inventory plots while shades of grey represent different inventory years - if applicable - with lighter shades representing earlier years.

**Figure 2:**
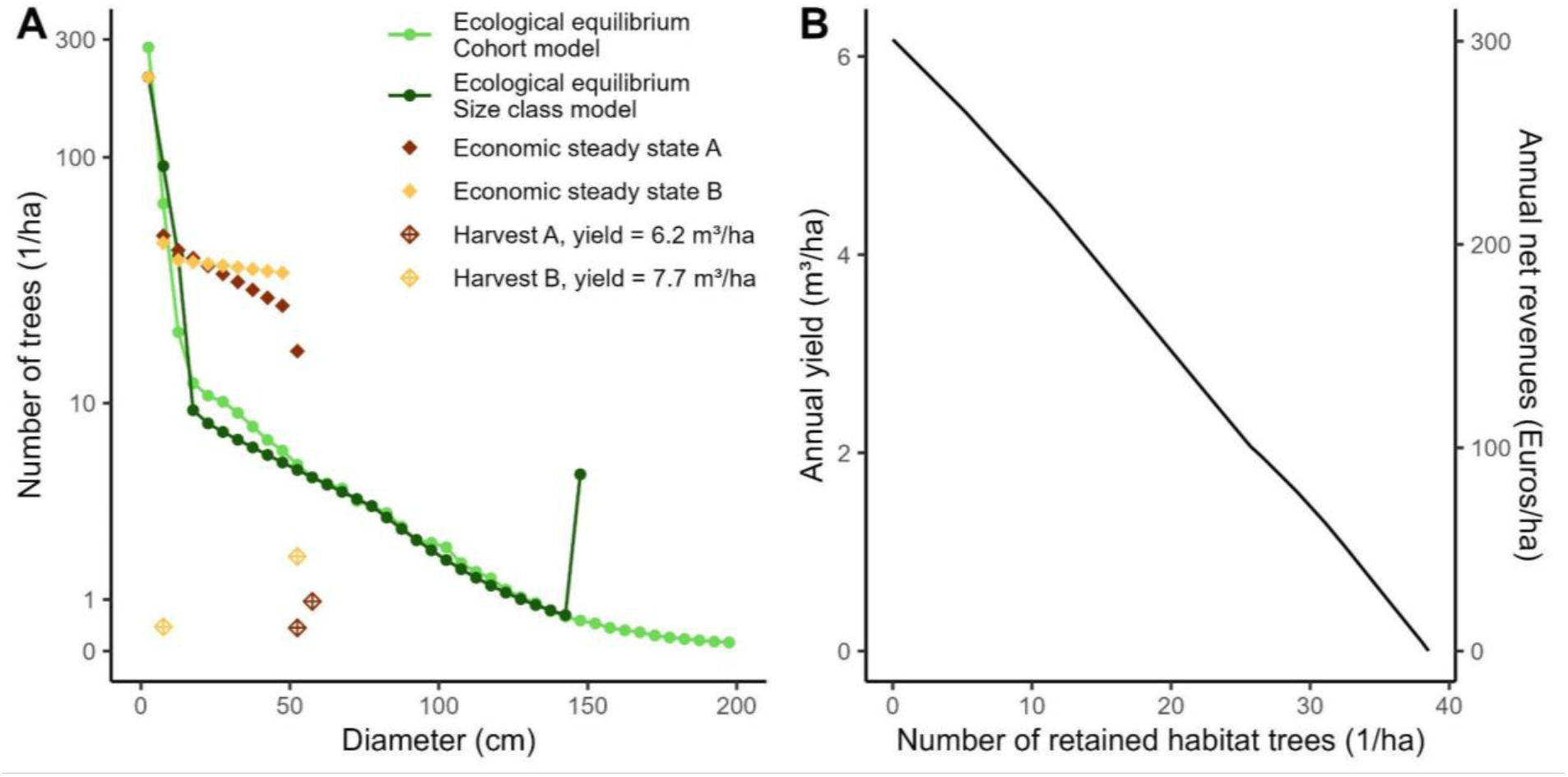
(A) Simulated size distributions in the ecological equilibrium of the cohort model (light green) and the size class model (dark green) along with the size distribution and optimal harvest in the economically optimal steady state using old-growth demographic rates (economic steady state A; red). Economic steady state B describes the same variables simulated with a canopy mortality rate of 0.1%. (B) Pareto frontier for the trade-off between timber yield (y-axis) and biodiversity conservation (as proxied by the number of retained habitat trees ≥70 cm dbh; x-axis).

### 3.2 Optimal harvesting

When maximising the net revenues from timber harvesting, the optimal management is to harvest all trees above a target diameter of 55–60 cm dbh. Net revenues are maximised when harvesting 0.44 trees of 55 cm dbh per hectare per year and all remaining trees larger than that (i.e., additional 0.96 trees of 60 cm dbh per hectare per year). This harvesting strategy amounts to a yield of 6.17 m^3^ of merchantable timber per hectare per year and hence to annual net revenues of approximately 300 Euros/ha. The remaining standing volume is 391 m^3^/ha at a basal area of 26.6 m^2^/ha and a density of 328 stems/ha (≥5 cm dbh). The remaining trees provide a total crown area of 1.28 m^2^/m^2^, such that only the trees in the smallest size class are in the understory. When reducing the canopy mortality rate *μ*_*C*_ to 0.1% the net-revenue-maximising management scenario entails the harvest of 0.46 trees of 10 cm dbh per hectare and year and a target dbh of 55 cm (2.00 trees/ha/y), amounting to a yield of 7.71 m^3^ and profits of 375 Euros per hectare per year. The remaining size distribution is flatter (**Figure 2A**), adding up to a standing volume of 393 m^3^/ha, a basal area of 27.0 m^2^/ha and a density of 333 stems/ha (≥5 cm dbh).

The maximum sustainable yield declines almost linearly with the number of retained habitat trees (≥70 cm dbh). With each additionally retained habitat tree, the maximum harvestable timber declines by about 0.16 m^3^/ha/y such that net revenues decline by 7.80 Euros/ha/y. Hence, at e.g., 10 habitat trees per hectare a maximum of 4.7 m^3^/ha/y timber can be harvested amounting to net revenues of 229 Euros/ha/y. The achievable yield drops to zero when retaining 38.5 or more habitat trees per hectare.

## 4. Discussion

We developed an ecological-economic optimization approach that combines ecological realism with mathematical tractability and, thus, the ability to be optimised numerically. This approach makes one of the best and widely tested inventory-calibrated forest simulation models available for rapid economic analyses. To achieve this, we transformed the cohort model into a size class model and parameterised both models for a beech-dominated forest in Central Germany. Both the cohort and the size class model reproduced ecological characteristics of unmanaged old-growth forests. The management strategy that maximises equilibrium net revenues in the size class model is similar to selective harvesting of beech forests practised and recommended in Germany (ThüringenForst/FFK Gotha, 2024). Our approach also offers a generalizable framework for the multi-criteria optimization of forests. Here, we focused on the trade-off between net revenues from timber harvesting and the retention of old habitat trees in equilibrium. Pareto frontiers quantifying this trade-off show an almost linear decline of maximum sustainable yield with each additional habitat tree, indicating that even moderate targets with respect to biodiversity conservation generate substantial costs.

The size class model approximated the cohort model well in terms of basal area and stem density of the forest in equilibrium. Even though both forest models are simple and use only three constant growth and mortality rates for trees in light, trees in the shade, and very large trees (>80 cm dbh), predictions of equilibrium forest structure of both models compared favourably to observed forest characteristics of unmanaged beech forests in Europe. Predicted basal area, density of large trees, and maximum diameter (for the cohort model) were at the upper boundary of observed values. There are three potential explanations for this result.

First, many of the currently unmanaged forests used for model validation have a management history, including the study site, and are still developing towards old-growth forests (Vandekerkhove et al., 2018). They are increasing over time in basal area, the number of large trees and maximum diameter (Figure 1). The predicted equilibrium forest structure is comparable to the structure of one of the few forests in Europe without management history (Uholka, Ukraine; Brändli et al., 2008). Second, both models use a constant mortality rate for very large trees (>80 cm dbh) while, in reality, mortality of larger trees increases strongly with diameter (Holzwarth et al., 2013). Finally, we only used trees for the calculation of demographic rates that were assigned to the two canopy layers with some certainty, i.e. trees that had a total crown area of larger trees of <80% or >120% of the plot area, respectively. This may also have led to a slight overestimation of canopy growth rates (but also to an overestimation of mortality rates because very large trees have higher mortality than mid-sized trees). That the cohort model is able to accurately project the structure and dynamics of unmanaged forests has previously been shown for temperate mixed-species forests in the US (Francis et al., 2023; Purves et al., 2008) and for tropical forests (Rüger et al., 2020). However, it was not clear whether this also holds for managed forests.

The cohort model shows periodic oscillations of 325 years at equilibrium (Appendix D, Fig. D1). Cohorts that recruit shortly before the simulated basal area (and hence crown area; Appendix D, Fig. D1B) reaches its minimum, become large enough for inclusion in the canopy just at this minimum point. Therefore, they spend most of their life in the canopy and grow very big due to the rather low canopy mortality. This results in an increase in basal area and crown area. As these cohorts begin to die off, a new decline in the stand basal area occurs, initiating a new cycle of oscillation. The size class model that we use for the economic evaluation, however, has a stable equilibrium solution so that these oscillations do not have consequences for the optimization.

The size class model identified the harvesting of trees when they reach a diameter of 55–60 cm as the management strategy that maximises net revenues from timber harvesting in equilibrium. This strategy is similar to selective harvesting of beech forests practised and recommended in Germany (Fritzlar and Biehl, 2006; Hessenmöller et al., 2012; Schütz, 2006; ThüringenForst, 2000; ThüringenForst/FFK Gotha, 2024). It is also in line with theoretical results for optimal equilibrium harvesting of age-structured populations (Reed, 1980). However, the predicted maximum yield is lower than yields in selectively managed beech forests in the same area (Hessenmöller et al., 2018, Schall et al., 2018). This is likely because we parameterized the model with demographic rates from unmanaged forests. While the effect of light availability on demographic rates is incorporated in the models through the distinction of two dynamic canopy layers, this may not fully capture the effect of targeted management interventions, including the promotion of target trees. We expect that in managed forests, mortality rates are lower than in unmanaged forests because slow-growing suppressed trees are harvested before they die (Meyer et al., 2022; Schall et al., 2018). This could also positively affect the growth rates of the remaining trees through a reduction of above- and below-ground competition that is not captured in the model. Average growth of trees >25 cm dbh in selectively harvested beech forests in the Hainich is ∼3 mm/y (Hessenmöller et al., 2018).

However, the difference between predicted and observed annual yield (6.2 versus 6.7-8.5 m^3^/ha) is small, which indicates that the models correctly capture the main ecological processes (Hessenmöller et al., 2012, 2018; Schall et al., 2018). Nevertheless, model predictions of maximum sustainable yield should be interpreted carefully. Moreover, the available allometric relationships for timber volume are uncertain for large trees (Hessenmöller et al., 2018). The fact that optimal forest management regulates forest density in a way that only the smallest trees (<10 cm dbh) are in the shaded understory, is also realistic (Hessenmöller et al., 2018). However, this result may also be caused by the fact that growth rates in the understory are likely to be underestimated and mortality rates overestimated, as only clearly assigned understory trees were included in the calculation of demographic rates. Another reason could be relatively low recruitment rates due to elevated browsing pressure at the Weberstedter Holz (Ohse et al., 2017; Schulze et al., 2014).

Predicted structural attributes of the remaining trees in equilibrium under optimal harvest (standing volume: 391 m^3^/ha, basal area: 26.6 m^2^/ha, stem density: 328 stems/ha) are similar to those observed in managed uneven-aged forests in Thuringia. While the Thuringian forestry authorities recommend a standing volume of up to 360 m^3^/ha (ThüringenForst, 2000; Fritzlar and Biehl, 2006), Hessenmöller et al. (2012) found an average standing volume of 401 m^3^/ha, a basal area of 27.1 m^2^/ha and a stem density of 352 stems/ha in forests in the Hainich area. The model of Hessenmöller et al. (2018) predicted a standing volume of up to ∼350 m^3^/ha at a basal area of 24 m^2^/ha and a stem density of ∼400 stems/ha, indicating the validity of our modelling and optimization approach.

We found the maximum sustainable yield to decrease approximately linearly with the retention of habitat trees for biodiversity conservation. Specifically, for each additional habitat tree retained per hectare, about 0.16 m^3^/ha/y of merchantable timber and 7.80 Euros/ha/y in net revenues are lost. The linearity of the trade-off can be attributed to the strict definition of the constraint for habitat trees, which can never be harvested but are left until their natural death. Since the optimal harvesting strategy targets diameters far below our threshold for habitat trees, the strategy itself does not change when retaining some trees as habitat trees, rather the number of harvested trees decreases linearly in consideration of the number of habitat trees required and of the natural mortality rate. Consequently, net revenues decrease proportionally. This result is in contrast to results for Boreal forests of Tahvonen et al. (2019), who found a concave trade-off between costs and a diversity index. However, their methodology significantly differed from our approach, which may account for the discrepancies.

Comparative values of lost revenues per retained habitat tree are scarce in the literature. Augustynczik et al. (2018) found reductions of 267 €/ha in the net present value of a forest stand over a 50 year management period when retaining 5 habitat trees per hectare (∼1.3 Euros/ha/y per habitat tree) in mixed montane forests in south-western Germany. Rosenkranz et al. (2014) find average losses of 0.4 - 0.66 m^3^/ha/y in timber harvest and 31 to 39 Euros/ha/y in profit when switching to conservational management regimes including the retention of habitat trees. However, Rosenkranz et al. (2014) do not provide the number of retained habitat trees. Assuming that the often recommended 3 to 10 habitat trees per hectare are retained and neglecting that the conservational management regime in Rosenkranz et al. (2014) includes further measures, such as the conservation of habitat-typical tree species, they find a range of 0.04 to 0.22 m^3^/ha/y or 3.1 to 13.0 Euros/ha/y per habitat tree, comparable to our results.

The payments for ecosystem services program “Klimaangepasstes Waldmanagement” (climate-adapted forest management) issued by the German Federal Ministry of Food and Agriculture promises forest managers compensations of 85 Euros/ha/y when adopting certain management criteria, among them retaining at least 5 habitat trees per hectare (BMEL, 2022). Although additional measures with potentially additional costs have to be taken, this seems to be an attractive program for forest owners considering lost net revenues of only 39 Euros/ha/a (5 × 7.80 Euros/ha/a).

We here developed a simple, generic forest model that can be coupled with effective numerical optimization, which is generalizable for any tree species. We expect the optimal management regime to depend on the ecological characteristics of the tree species. European beech is a very shade-tolerant tree species, and saplings can remain in the shaded understory for decades and wait for a canopy gap to be able to grow up into the canopy (Feldmann et al., 2018; Stiers et al., 2019). While we plan to analyse this in future studies, we suspect that, for light-demanding tree species, understory mortality might be much higher and natural recruitment under a shading canopy might be rare or absent. For species with these characteristics, optimal management would have to keep only one canopy layer to allow for successful natural regeneration. If forest management would be optimised over time and tree planting (artificial regeneration) would be included, clear-cut harvesting would be another candidate for optimal management.

This modelling framework is a major improvement for the mathematical optimization of forest management that, in Central Europe, has currently only been carried out for single-species or even-aged forests (Assmuth et al., 2018). We expect that the framework can easily be extended to mixed-species forests, where a mathematical optimization has only been performed for boreal forests (Tahvonen et al., 2019).

While this study is a first step towards a general optimization framework for the management of temperate forests, we suggest several lines of improvement for future work. From the theoretical perspective, dynamic, size-dependent harvesting costs and wood prices should be included to identify economically feasible management options. This will increase the results’ level of realism and ensure that the needs of the timber market are met. Maximum profits might be achieved with less timber harvest if costs were negatively correlated with size, and, hence, the trade-off with the number of retained habitat trees might be shaped differently. Moreover, in addition to finding the equilibrium solution for optimal management, it is also necessary to be able to optimise harvesting strategies over time. This way, favourable management scenarios can be identified also for the transition towards the economically optimal equilibrium. We expect that discounting also may have a major effect on the optimal harvesting strategy and the trade-off between revenues from timber harvesting and biodiversity maintenance. For such a sensitivity analysis of the optimal harvesting strategy, an analytical approach to solving the ecological-economic optimization model would be very useful. The model we present here might serve as an excellent starting point for such an analysis in future work. From the applied perspective, the extension of the ecological-economic optimization framework to mixed-species forests will be the next step. Researchers and forest management policies demand the conversion of monocultures to mixed-species forests to increase the resilience of forests to climate change (Gamfeldt et al., 2013; Knoke et al., 2008; Schuler et al., 2017). Moreover, ecosystem services that can be quantified in monetary terms, are straightforward to include in the optimization model. Incorporating additional ecosystem services, such as carbon storage, climate resilience, or, in densely-populated areas, the recreational value, is of increasing relevance in the face of global change.

## Acknowledgements

We thank Jürgen Huss for establishing the large-scale inventory site ‘Weberstedter Holz’ in 1999 and his successor Jürgen Bauhus for supporting the re-inventory in 2007 and his collaboration. We thank all field workers who collected the forest inventory data and the Max-Planck-Society and the German Centre for Integrative Biodiversity Research (iDiv) Halle-Jena-Leipzig (DFG, FZT 118) for funding the second and third forest inventories. We are grateful to the administration of the Hainich National Park for the fruitful collaboration. We thank Peter Schall, Martina Mund and Felix Meier for helpful discussions and comments. Stefan Fleck and Fréderic Holzwarth kindly provided tree crown area and height data. NR acknowledges funding from the Senior Scientist Programme of iDiv (FZT 118).

## Author contributions

**Markus E. Schorn:** Software, Validation, Formal analysis, Writing - Original Draft, Writing - Review & Editing, Visualization. **Martin F. Quaas:** Conceptualization, Methodology, Software, Writing - Review & Editing, Supervision, Funding acquisition. Hanna Schenk: Software, Validation, Formal analysis,Writing - Review & Editing. **Christian Wirth:** Writing - Review & Editing. **Nadja Rüger:** Conceptualization, Methodology, Writing - Original Draft, Writing - Review & Editing, Supervision, Funding acquisition.

## Declaration of Generative AI and AI-assisted technologies in the writing process

During the preparation of this work the authors used ChatGPT in order to improve the language of some paragraphs. After using this tool/service, the authors reviewed and edited the content as needed and take full responsibility for the content of the publication.

