## Supplementary Material for "Multi-objective management of naturally regenerating beech forests – An ecological-economic optimization approach"

**Appendix A**

**Model parameters α and β**

The parameters α and β in the size class model describe the rate at which trees are promoted from one size class to the next larger size class within one theoretical time step (α) or at which trees remain in the same size class (β). Both depend on the growth and mortality rates derived from forest inventory data. They are defined for each size class. α and β differ for understory and canopy size classes and for size classes that are 80 cm or larger, i.e., they also depend dynamically on the shading from larger trees to which the respective size class is exposed. Further, if the size class is 80 cm dbh or larger, α and β of that size class are $\alpha_L$ and $\beta_L$. If the shading from above is smaller than 1, the size class is in the canopy, such that α and β of that size class approximate $\alpha_C$ and $\beta_C$. If the shading from above is smaller than 1, α and β approximate $\alpha_U$ and $\beta_U$. The transition from $\alpha_U$ and $\beta_U$ to $\alpha_C$ and $\beta_C$ is defined in Equations 3 and 4 in the main manuscript and is shaped by a smoothing factor σ = 0.01. The transition of $\alpha_U$ to $\alpha_C$ is depicted in Figure A1 as an example. $\alpha_{U,}$ $\alpha_{C,}$ $\alpha_{L,}$ $\beta_{U,}$ $\beta_{C,}$ and $\beta_L$ are defined in Equations 1 and 2 in the main manuscript.

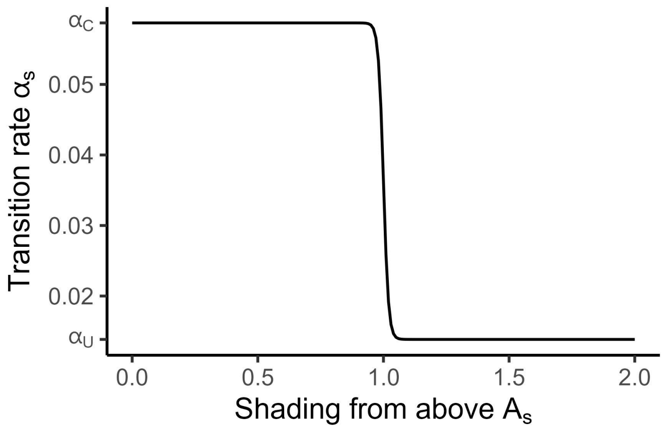


**Figure A1:** Transition of $\alpha_s$ of a size class s from approximating $\alpha_C$ to approximating $\alpha_U$. The transition is shaped by the smoothing factor σ = 0.01 and happens when the shading from larger trees surpasses 1, such that the size class is fully shaded and, hence, in the understory.

**Appendix B**

**Canopy layer classification**

To assign trees to the canopy or the understory, we estimate height and crown area of each tree from its diameter at breast height (dbh) using species-specific allometric equations. We estimated these allometric relationships based on data on height, crown projected area and dbh from 1,316 trees in the Weberstedter Holz (Fleck et al., 2011; Holzwarth et al., 2015).

For beech (*Fagus sylvatica*) and hornbeam (*Carpinus betulus*), we fit species-specific relationships based on data from 842 and 35 trees, respectively. For lime (*Tilia* spp.) and maple (*Acer* spp.), we fit genus-specific relationships based on data from 366 and 73 trees, respectively (Tables B1 & B2, Figures B1 & B2). We used the parameters for beech (*Fagus sylvatica*) also to estimate height and crown areas of oak (*Quercus robur*) and ash (*Fraxinus excelsior*) trees and the parameters for lime (*Tilia* spp.) to estimate height and crown areas of all other species. We fit the relationships using the *least_squares* function from the *scipy.optimize* package for *python*.

**Diameter-crown area relationships**

Allometric equations for dbh - crown area relationships were estimated based on the form:

$$ca = \frac{a}{1+exp(\frac{c-dbh}{d})} + \frac{b}{1+exp(\frac{c-dbh}{d})}.$$

Dbh is in cm, crown area (ca) is in m². Model parameters are given in Table B1 and the model fit is shown in Figure B1.

**Table B1**: Model parameters for the dbh - crown area allometric relationship in the Weberstedter Holz for beech, hornbeam, lime and maple.

| **Species** | **a** | **b** | **c** | **d** |
|---|---|---|---|---|
| *Fagus sylvatica* | 10.31 | 250 | 18.06 | 77.06 |
| *Carpinus betulus* | -29.13 | 200 | 26.56 | 46.54 |
| *Tilia* spp. | -0.50 | 200 | 19.00 | 64.93 |
| *Acer* spp. | -0.40 | 200 | 22.30 | 65.58 |

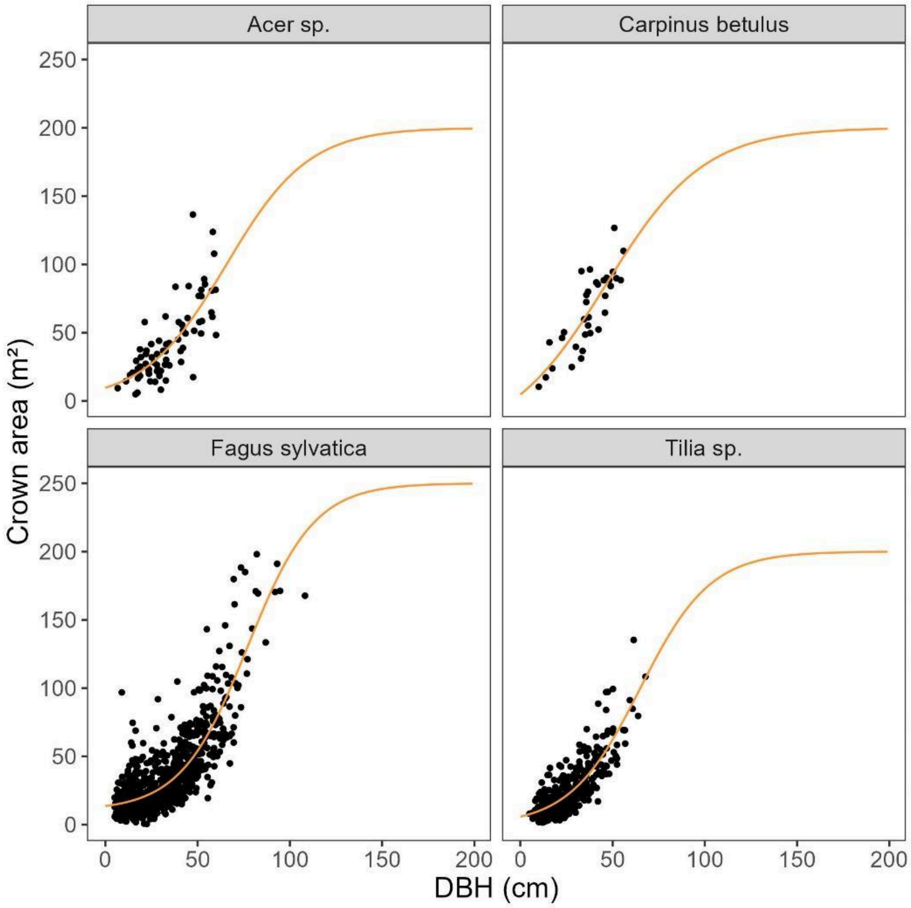


**Figure B1**: Data (black dots) and derived models (orange lines) on dbh - crown area allometric relationships for species (or genuses) from the Weberstedter Holz. The model parameters are given in Table B1.

**Diameter - height relationships:**

Allometric equations for dbh - height relationships were estimated based on the form (adapted from Mund 2004):

$$H = 1.3 + \left(\frac{dbh}{a + b \cdot dbh}\right)^3$$

Dbh is in cm, height (H) is in m. Model parameters are given in Table B2 and the model fit is shown in Figure B2.

**Table B2**: Model parameters for the dbh - height allometric relationship in the Weberstedter Holz for beech, hornbeam, lime and maple.

| **Species** | **a** | **b** |
|---|---|---|
| *Fagus sylvatica* | 1.649 | 0.285 |
| *Carpinus betulus* | 1.538 | 0.311 |
| *Tilia* spp. | 1.354 | 0.306 |
| *Acer* spp. | 1.564 | 0.304 |

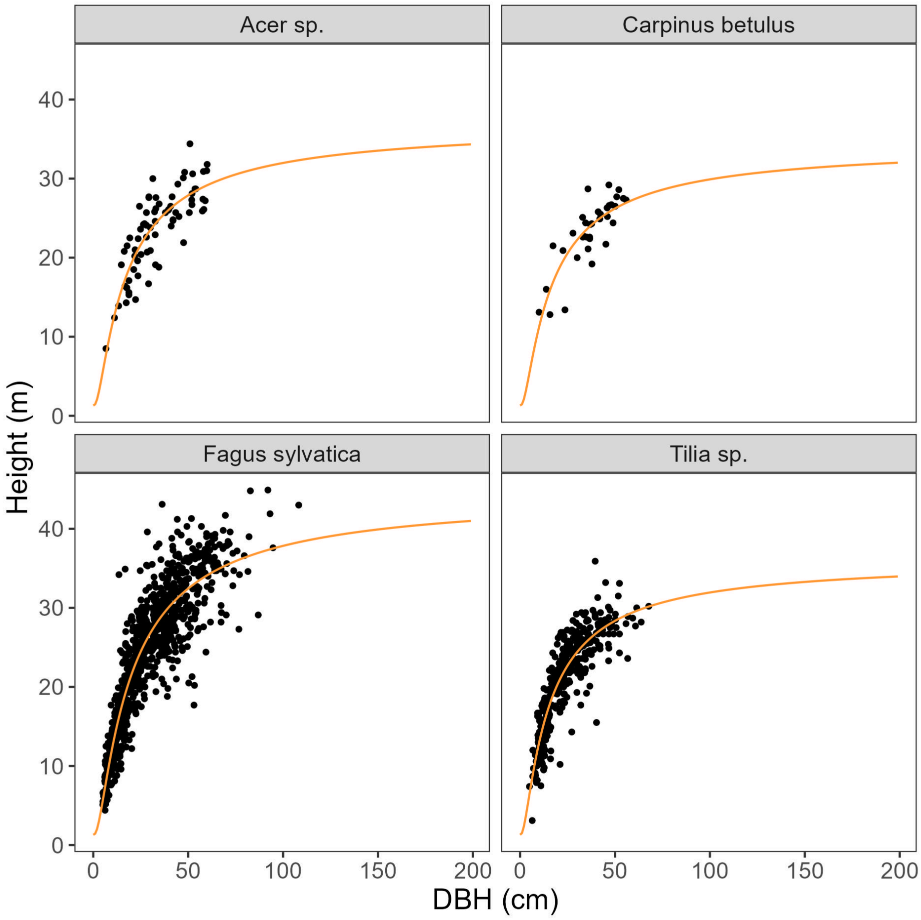


**Figure B2**: Data (black dots) and derived models (orange lines) on dbh - height allometric relationships for species (or genuses) from the Weberstedter Holz. The model parameters are given in Table B2.

**Appendix C**

**Recruitment rates**

We calculated the number of individuals that surpassed the 1.3 m height threshold per year and ha as well as shading (total crown area/400 m²) for each of the 400-m² subplots in both census intervals. We then calculated the mean recruitment across all subplots with shading <1 or ≥1 and used these as open-canopy or closed-canopy recruitment rates, respectively (Figure C1).

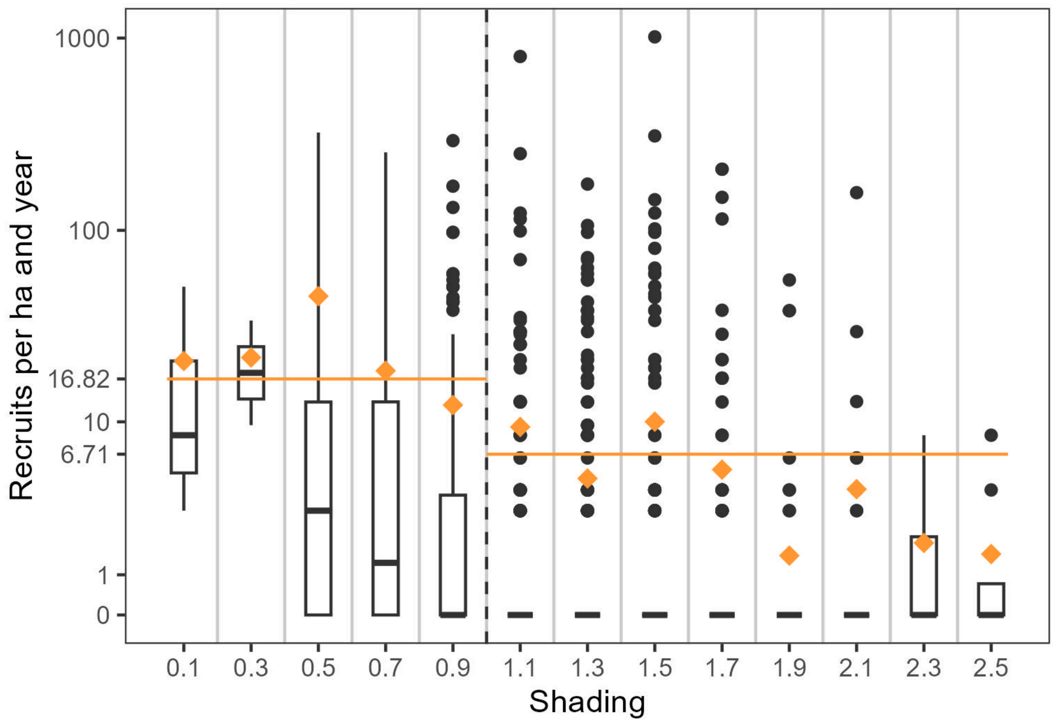


**Figure C1:** Boxplots for the number of recruits per ha and year for each of the 400 m² subplots grouped by total shading. Orange diamonds represent the mean per shading class and orange lines represent the mean number of recruits in open-canopy (shading<1) and closed-canopy (shading≥1) conditions.

**Appendix D**

**Simulations over time of the cohort model**

As described in the main manuscript, the cohort model performs regular oscillations of a length of 325 years at equilibrium (see Figure D1). All structural attributes, that are used to validate the model against the size class model and values from unmanaged beech forests in Europe, are therefore averaged across one oscillation cycle. Since we model a maximum of 2000 years, all values are the mean of the years 1675 - 2000.

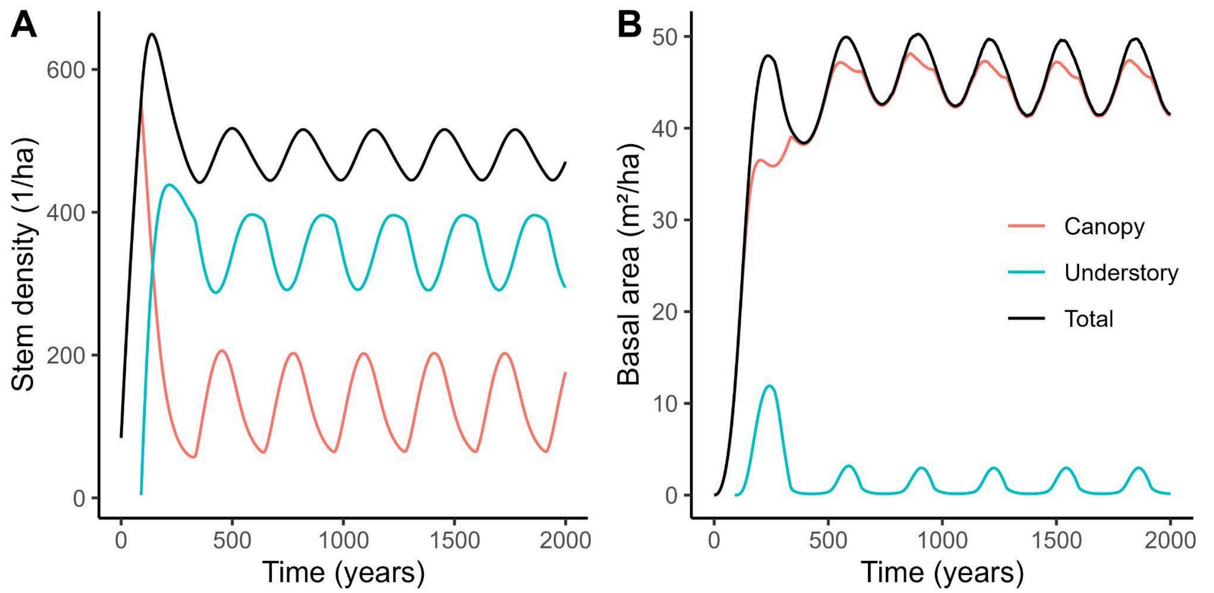


**Figure D1:** Canopy, understory and total stem density and basal area over time as modeled with the cohort model starting from bare ground.

**Appendix E**

**Validation**

Model outputs were validated against reference values from available data or literature values from comparable old-growth beech forests in Central and Eastern Europe (Table E1).

Table E1: Basal area of all trees ≥30 cm dbh, density of very large trees (VLT, ≥80 cm dbh) and maximum dbh for the models and reference values from available data and the literature.

| **Source** | **Forest inventory plot** | **Year** | **Basal area ≥30 cm dbh ($m^2$/ha)** | **Density VLT (n/ha)** | **Dbh max (cm)** | **Above-ground biomass (t/ha)** | **Stem density ≥5 cm dbh (n/ha)** |
|---|---|---|---|---|---|---|---|
| Size class model | | | 41.9 | 30.1 | 150 | 530 | 235 |
| Cohort model | | | 43.4 | 27.2 | 170.5 | 556 | 201 |
| Data | Weberstedter Holz | 1999 | 31.9 | 6.8 | 126.5 | 362 | 383 |
| | | 2007 | 33.4 | 9.3 | 135.5 | 380 | 359 |
| | | 2013 | 35.8 | 11.6 | 147.7 | 408 | 364 |
| Stillhard et al. (2019) | Uholka (core area) | 2000 | 40.5 | 23.3 | 132 | 494 | 289 |
| | | 2005 | 42.3 | 27.5 | 133.2 | 520 | 302 |
| | | 2010 | 42.5 | 28.8 | 133.2 | 527 | 328 |
| | | 2015 | 40.6 | 29.9 | 129 | 510 | 433 |
| Vandekerkhove et al. (2018) | Heilige Hallen | 2000 | 24.4 | 19.0 | 148 | | |
| | Limker Strang | 2000 | 30.2 | 0.7 | 93 | | |
| | Mirdita | 2000 | 37.2 | 5.4 | 99 | | |
| | Puka | 2000 | 45.4 | 15.4 | 122 | | |
| | Rajka | 2000 | 43.4 | 19.3 | 115 | | |
| | Razula | 1972 | 25.7 | 4.5 | 125 | | |
| | | 1995 | 26.8 | 10.3 | 129 | | |
| | | 2009 | 27 | 12.4 | 146 | | |
| | Salajka | 1974 | 26.2 | 19.3 | 190 | | |
| | | 1994 | 24.3 | 14.2 | 142 | | |
| | | 2007 | 27 | 14.3 | 135 | | |
| | Uholka (sampling plots) | 2010 | | 12.0 | 150 | | |
| | Zofin | 1975 | 35.1 | 26.0 | 140 | | |
| | | 1997 | 34.3 | 23.0 | 145 | | |
| | | 2008 | 30.4 | 21.2 | 146 | | |
| | Kersselaerspleyn | 1986 | 28.4 | 31.3 | 135 | | |
| | | 2001 | 30.6 | 33.5 | 150 | | |
| | | 2011 | 31.3 | 34.3 | 158 | | |

### Literaturverzeichnis

Fleck, S., Mölder, I., Jacob, M., Gebauer, T., Jungkunst, H.F., Leuschner, C., 2011. Comparison of conventional eight-point crown projections with LIDAR-based virtual crown projections in a temperate old-growth forest. Ann. For. Sci. 68, 1173–1185. DOI: 10.1007/s13595-011-0067-1

Holzwarth, Frédéric; Rüger, Nadja; Wirth, Christian (2015): Taking a closer look: disentangling effects of functional diversity on ecosystem functions with a trait-based model across hierarchy and time. In: *Royal Society open science* 2 (3), S. 140541. DOI: 10.1098/rsos.140541.

Mund, Martina (2004): Carbon pools of European beech forests (Fagus sylvatica) under different silvicultural management. PhD-Thesis. Georg-August-Universität, Göttingen.

Stillhard, Jonas, Martina Hobi, Lisa Hülsmann, Peter Brang, Christian Ginzler, Myroslav Kabal, Jens Nitzsche, Gilbert Projer, Yuriy Shparyk, and Brigitte Commarmot (2019): Stand Inventory Data from the 10-Ha Forest Research Plot in Uholka: 15 Yr of Primeval Beech Forest Development. In: *Ecology* 100 (11): e02845. DOI: 10.1002/ecy.2845.

Vandekerkhove, Kris; Vanhellemont, Margot; Vrška, Tomáš; Meyer, Peter; Tabaku, Vath; Thomaes, Arno et al. (2018): Very large trees in a lowland old-growth beech (Fagus sylvatica L.) forest: Density, size, growth and spatial patterns in comparison to reference sites in Europe. In: *Forest Ecology and Management* 417, S. 1–17. DOI: 10.1016/j.foreco.2018.02.033.
